# Validity and reliability of Kinect v2 for quantifying upper body kinematics during seated reaching

**DOI:** 10.1101/2022.01.18.476737

**Authors:** Germain Faity, Denis Mottet, Jérôme Froger

## Abstract

**Background:** Kinematic analysis of the upper limbs is a good way to assess and monitor recovery in individuals with stroke, but remains little used in clinical routine due to its low feasibility. The aim of this study is to assess the validity and reliability of the Kinect v2 for the assessment of 17 kinematic variables commonly used in the analysis of upper limb reaching in stroke.

**Methods:** 26 healthy participants performed seated hand-reaching tasks while holding a dumbbell to induce a behaviour similar to that of a person with a stroke. 3D upper limb and trunk motion were simultaneously recorded with the Kinect v2 (Microsoft, USA) and with the VICON (OxfordMetrics, UK), the latter being the reference system. For each kinematic outcome, the validity of the Kinect was assessed with ICC, linear regression and Bland & Altman plots.

**Results:** The Kinect assesses trunk compensations, hand range of motion, movement time and mean velocity with a moderate to excellent reliability. In contrast, elbow and shoulder range of motion, time to peak velocity and path length ratio have a poor to moderate reliability, indicating that these variables should be interpreted with caution. Finally, instantaneous hand and elbow tracking are not precise enough to reliably assess Cartesian and angular kinematics over time, rendering variables such as the number of velocity peaks and the peak hand velocity unusable.

**Conclusions:** Thanks to its ease of use and markerless properties, the Kinect can be used in clinical routine for semi-automated quantitative diagnostics guiding individualised rehabilitation of the upper limb. However, engineers and therapists must bear in mind the limitations of the Kinect for the instantaneous tracking of the hand and elbow.

## 1) Background

Stroke results in major movement deficits, especially in the upper limb. To provide the best possible rehabilitation, therapists regularly assess the motor performance of individuals with stroke. However, the clinical scales used by therapists suffer from several limitations. First, clinical scales have a subjective scoring system which limits the reliability of ratings between therapists and over time [1]. Second, clinical scales are too often insensitive to changes. For example, in most of its items, the Fugl-Meyer Assessment only supports 3 rating levels (0, 1 or 2), whereas it is considered as one of the strongest clinical scales [2]. Third, most clinical scales do not sufficiently account for compensations that may occur in stroke movement [3, 4], which cancels out the differences between a true recovery and a compensation pattern [5].

To go beyond these limitations, scientists use motion capture to quantify the motor deficits. Indeed, upper limb and trunk kinematics are more sensitive to changes than clinical scales [6, 7], and can even predict motor outcomes over several months [8, 9]. Moreover, motion capture makes compensation assessment easy and more objective [3, 10, 11]. Despite these advantages, kinematic assessment of the upper limb remains little used in clinical practice because of its poor feasibility. Indeed, motion capture systems are expensive and require a large volume to perform the movements. In addition, patients have to be suited up with markers placed with accuracy, which takes precious time in the clinical context. Finally, these technologies require a high technical level to extract valuable variables from raw data.

Since its release in 2013, the Kinect v2 (also known as Kinect One or Kinect for Xbox One, Microsoft, USA) has been widely used for rehabilitation purpose and has largely contributed to the rise of virtual reality in rehabilitation trials. Virtual reality with Kinect may be beneficial in improving upper limb function and activities of daily living when used as an adjunct to usual care (to increase overall therapy time) [12, 13]. However, although the markerless and ease of use properties of the Kinect v2 facilitate its use in clinical routine, and its value for gait analysis has been documented [14], the validity of the Kinect for assessing upper limb kinematics after stroke remains to be tested.

Previous works shows that the Kinect v2 has an average accuracy of 10-15 mm but can generate distance errors up to 80 mm [15, 16]. In addition, the Kinect would detect range of motion (ROM) with 1 to 10° error [17–20] but this result should be taken carefully given the variability between the studies. Indeed, some authors argue that the Kinect v2 has excellent reliability, especially for flexion of the elbow and shoulder [21], but others conclude that the kinematics obtained by the Kinect are unreliable [22] and that the use of IMUs should be preferred for motor assessment [23].

In people with stroke, when assessed by the Kinect v2, hand and trunk range of motion are valid and reliable [24] and kinematics can distinguish the reaching performance between healthy control, less-affected side and more-affected side of patients with stroke [25–27]. Yet, there is no further information on precise kinematic variables.

**The goal of our study is to investigate to what extent the Kinect v2 is valid and reliable for the kinematic assessment of reaching movements in people with stroke**. To do so, we simultaneously recorded reaching movements with the Kinect v2 and the Vicon motion capture system considered as gold-standard. We hypothesized that the Kinect v2 will provide the same information as the Vicon system.

Because it was not possible to ask patients to come to the laboratory, we tested the reliability of the Kinect v2 with a model of stroke behaviour [28]. Specifically, we asked healthy participants to perform a series of reaching movements with their hand loaded to 75% of their maximum voluntary antigravity torque. In this loaded condition, healthy participants spontaneously develop compensations similar to those observed in most stroke patients, including trunk flexion and rotation, reduced shoulder abduction, and reduced movement speed.

## 2) Methods

### 2.1. Participants

26 healthy participants (12 males, age 21 ± 3 years, 3 left-handed, height 1.73 ± 0.09 m, weight 66.92 ± 9.29 kg) took part to this study.

Participants were not included if they had shoulder pain or any other problem that could affect their movement. This study was performed in accordance with the 1964 Declaration of Helsinki. The local ethics committee approved the study (IRB-EM 1901C).

### 2.2. Experimental protocol

Participants performed the PANU test [3] as described in a previous study [28]. Participants had to reach a target with the side of their thumb nail. The target was a table tennis ball fixed in front of the participant at a height of 0.80 m, just within the anatomical reaching distance for the hand. The starting position was seated, feet on the ground, back in contact with the chair and forearms on the armrests.

Participants completed 5 reaches both in the spontaneous trunk use condition and in the restrained trunk use condition. In the spontaneous trunk use condition, participants had to reach the target at a natural pace, wait 1 second and return to the starting position. In the restrained trunk use condition, participants had to reach the target while minimising trunk movement: the experimenter manually applied a light proprioceptive feedback on the participant’s shoulders, as a reminder to minimise trunk movement. We did not use a belt to restrain the trunk in order to leave the participant free to use the trunk if necessary, and thus avoid task failure.

The assessed hand was chosen pseudo-randomly (12 left, 14 right) so that half of the participants performed the task with their dominant hand, and the other half with their non-dominant hand. The weight of the arm including the dumbbell was set to 75.0% (± 5.5%) of the maximum antigravity force (MAF) in the posture with the hand at the target.

### 2.3. Experimental setup

The movements of the participants were recorded by both a Vicon motion capture system and a Kinect v2. The data obtained by the Kinect were then compared to the data obtained by the Vicon, the latter being considered as the ground truth.

#### 2.3.1. Vicon sensor

The Vicon system (Oxford Metrics, U.K.) is a marker-based optoelectronic motion capture tool that is widely used for kinematic measurements [29]. Indeed, with a similar setting to ours, the error of the Vicon is 0.15 mm ± 0.025 mm in static and remains less than 2 mm in dynamic [30]. In this study, we used a 6-camera rectangle Vicon system with a sampling frequency of 100 Hz. Vicon time series were recorded using “Vicon Nexus 2” software (Oxford Metrics, U.K.).

#### 2.3.2. Kinect sensor

The Kinect v2 (Microsoft, USA) is a markerless motion capture tool combining 3 sensors (a RGB colour camera, a depth sensor and an infrared sensor) to provide the 3D position of 25 landmarks on a skeleton with a sampling rate of 30 Hz [24]. The Kinect was connected to a PC running the “MaCoKi” software (NaturalPad, Montpellier, France) developed from the Kinect SDK (v2.0_1409, Microsoft, USA) to record the position time-series of the hands and trunk. As recommended in previous studies, we placed the Kinect in front of the participant, at a distance of 1.50 m, a height of 1.40 m and with no direct sunlight to minimise errors [18, 31–33] (Fig. 1).

**Figure 1.**
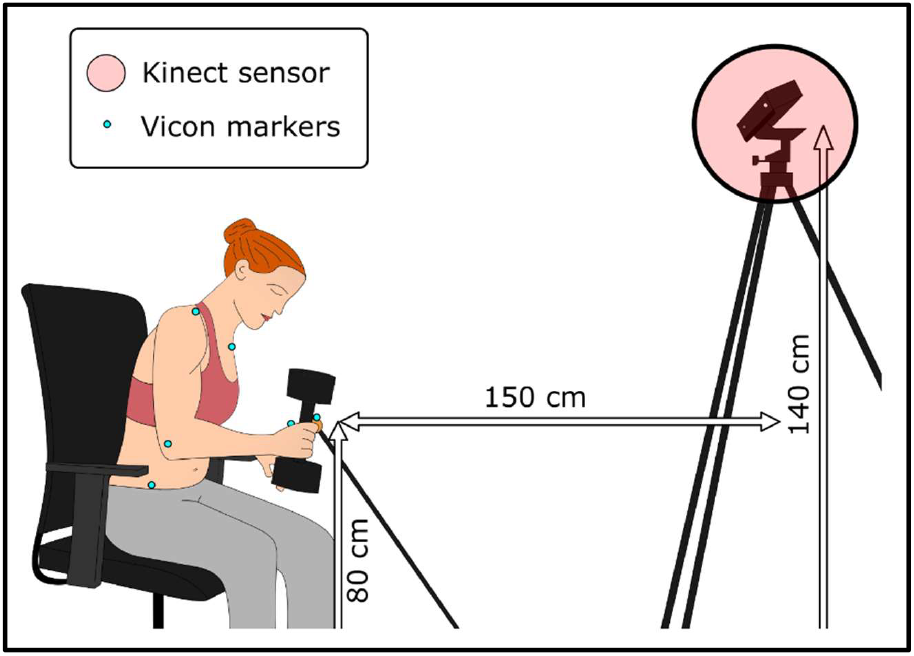
Experimental setup. 3D upper limb kinematics were simultaneously recorded by the Kinect and the Vicon motion capture systems. Vicon markers were placed on the target, hands, elbows, shoulders, manubrium and hips of the participant as close as possible to the joint centres located by the Kinect. The Kinect was located in front of the participant, at a distance of 1.50 m and a height of 1.40 m. The target (orange table tennis ball) was located at a height of 0.80 m, just within the anatomical reaching distance for the hand.

#### 2.3.3. Position of landmarks for the Vicon and for the Kinect

In order to compare the Kinect data to the Vicon data, we placed the Vicon markers as close as possible to the joint centers located by the Kinect. Thus, we placed markers at the manubrium (spine-shoulder for Kinect), and for each body side on the 1st metacarpal (wrist for Kinect), the lateral epicondyle of the humerus (elbow for Kinect), the acromion process (shoulder for Kinect), and the anteriosuperior iliac spine. For each side, we corrected the anteriosuperior iliac spine marker position before data analysis to best match the anatomical center of hips joints (hips for Kinect). In order to facilitate the reading, the spine shoulder marker is renamed “trunk marker” in the text.

### 2.4. Data processing

Data processing was performed with SciLab 6.0.2. The Kinect being positioned obliquely to the ground to minimise visual occlusions of the elbow, we realigned the Kinect axes to match the Vicon axes. This realignment was performed using a solid transformation of the Kinect data based on the “Least-Squares Fitting” method [34] implemented in a matlab function by Nghia Ho [35].

All position time series were low pass filtered at 2.5 Hz with a dual pass second order Butterworth filter. We choose a cut-off frequency of 2.5 Hz because the analysis of raw data showed that the frequency band 0-2.5 Hz contains at least 95% of the spectral density of the time series data.

We first calculated the start and end of each reaching movement in the one-dimensional task space [28]. Because the goal of a reaching task is to bring the hand to the target, that is, to reduce the hand-to-target distance, what is important for task success is the hand-to-target Euclidean distance. The hand-to-target Euclidean distance summarises the 3D effector space into a 1D task space (where movement matters) leaving aside a 2D null space (where movement does not impact task success). We fixed the beginning of the movement (t0) when the Euclidean velocity of the hand in task space became positive and remained positive until the maximum velocity. The end of the movement (tfinal) was when the Euclidean distance to the target reached its minimum.

Angles presented in this study were calculated as the difference between the anatomical angle at tfinal and the anatomical angle at t0, as described in the equation 1.

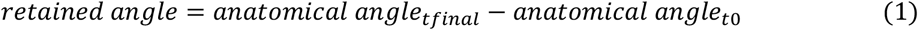

### 2.5. Statistical analysis

We assessed the degree of reliability between the Kinect and the Vicon variables using intraclass coefficient correlation (ICC) and coefficient of determination (r^2^). We complemented these measures with Bland and Altman plots to evaluate the validity of the Kinect through the difference in means and to estimate an agreement interval through the 95% limits of agreement [36]. To compare validity across variables, a relative systematic error was calculated as 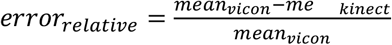

ICC estimates were calculated using R (version 3.6.1) based on a single-rating, consistency, one-way random-effect model. As stated by Koo & Li, *“values less than 0.5 are indicative of poor reliability, values between 0.5 and 0.75 indicate moderate reliability, values between 0.75 and 0.9 indicate good reliability, and values greater than 0.90 indicate excellent reliability”* [37]. We used the same limits for the *error*_relative_. The level of significance for all tests was set at p < .05. All coefficients of determination r^2^ were found to be statistically significant.

## 3) Results

The results are reported in table 1.

**Table 1.**
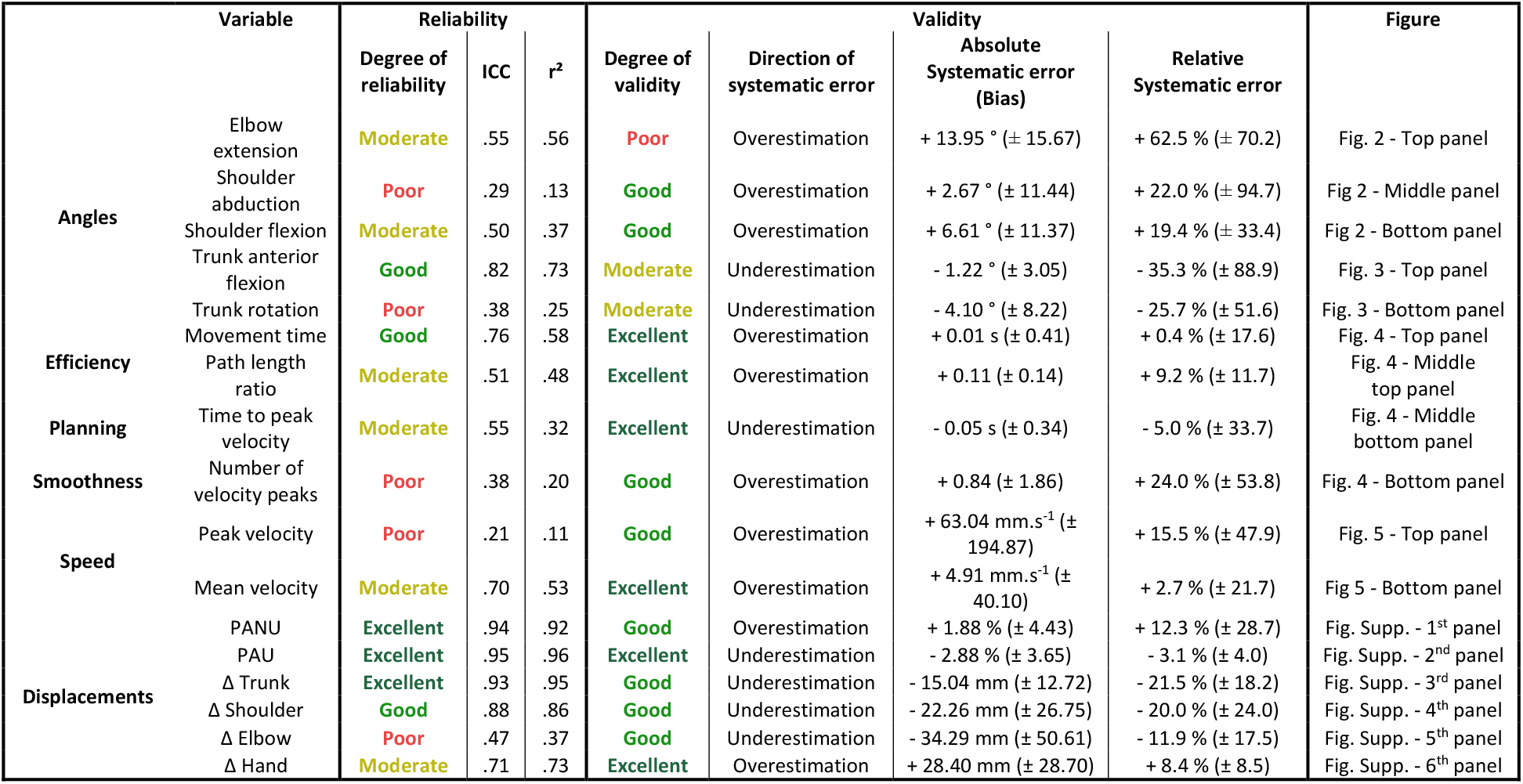
Reliability and validity of the Kinect main kinematic variables used in the analysis of reaching in stroke [38]. Reliability measures the consistency of the results compared to the ground truth (i.e., for each individual, how close the Kinect measure is to the ground truth). A perfect reliability between Kinect and Vicon data would result in an intraclass correlation coefficient (ICC) of 1 and a coefficient of determination (r^2^) of 1 as well. The reliability of each of the main variables is summarised on the Y axis of the figure 6 using the ICC. Validity measures the extent to which the results are close to the ground truth on average (i.e., the higher the percentage of error on average, the lower the validity). A perfect validity would result in a difference in means of 0 in the Bland and Altman plot, and thus in a relative systematic error of 0. The validity of each of the main variables is summarised on the X axis of the figure 6 using the relative systematic error.

### 3.1. Angles

**Figure 2.**
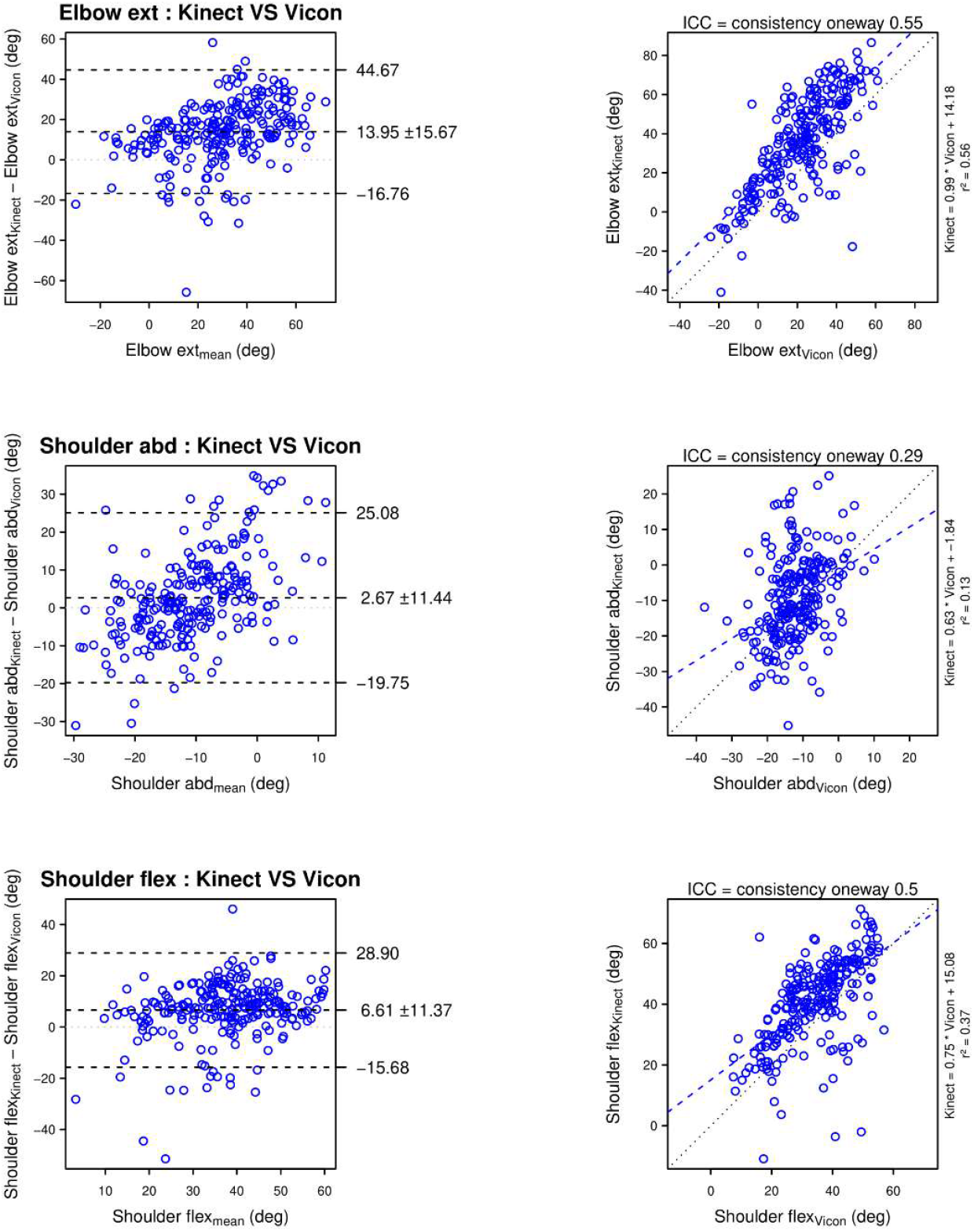
Comparison of the arm and shoulder angles assessed with the Kinect and the Vicon systems. Panels in the first row illustrate elbow extension (Elbow ext). Panels in the second row illustrate shoulder abduction (Shoulder abd). Panels in the third row illustrate shoulder flexion (Shoulder flex). For each row, the left panel represents the Bland and Altman plot and the right panel represents the linear regression plot. When assessed with the Kinect, elbow extension is moderately reliable but highly overestimated, shoulder abduction is poorly reliable and slightly overestimated in average, shoulder flexion is moderately reliable and slightly overestimated.

**Figure 3.**
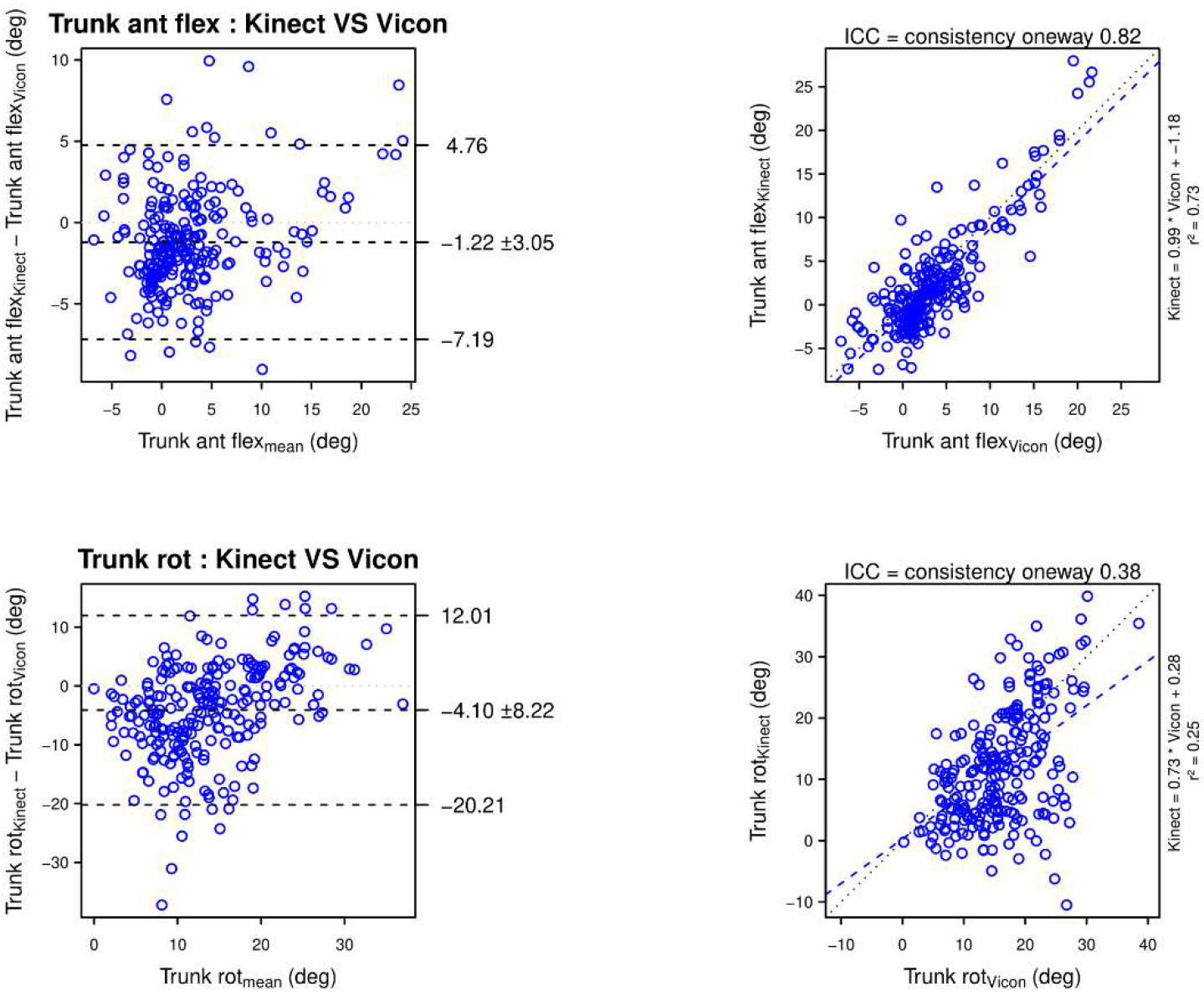
Comparison of the trunk angles assessed with the Kinect and the Vicon systems. Panels in the first row illustrate trunk anterior flexion (Trunk ant flex). Panels in the second row illustrate trunk rotation (Trunk rot). For each row, the left panel represents the Bland and Altman plot and the right panel represents the linear regression plot. When assessed with the Kinect, trunk anterior flexion is goodly reliable but moderately underestimated, trunk rotation is poorly reliable and moderately underestimated.

### 3.3. Efficiency, planning and smoothness

**Figure 4.**
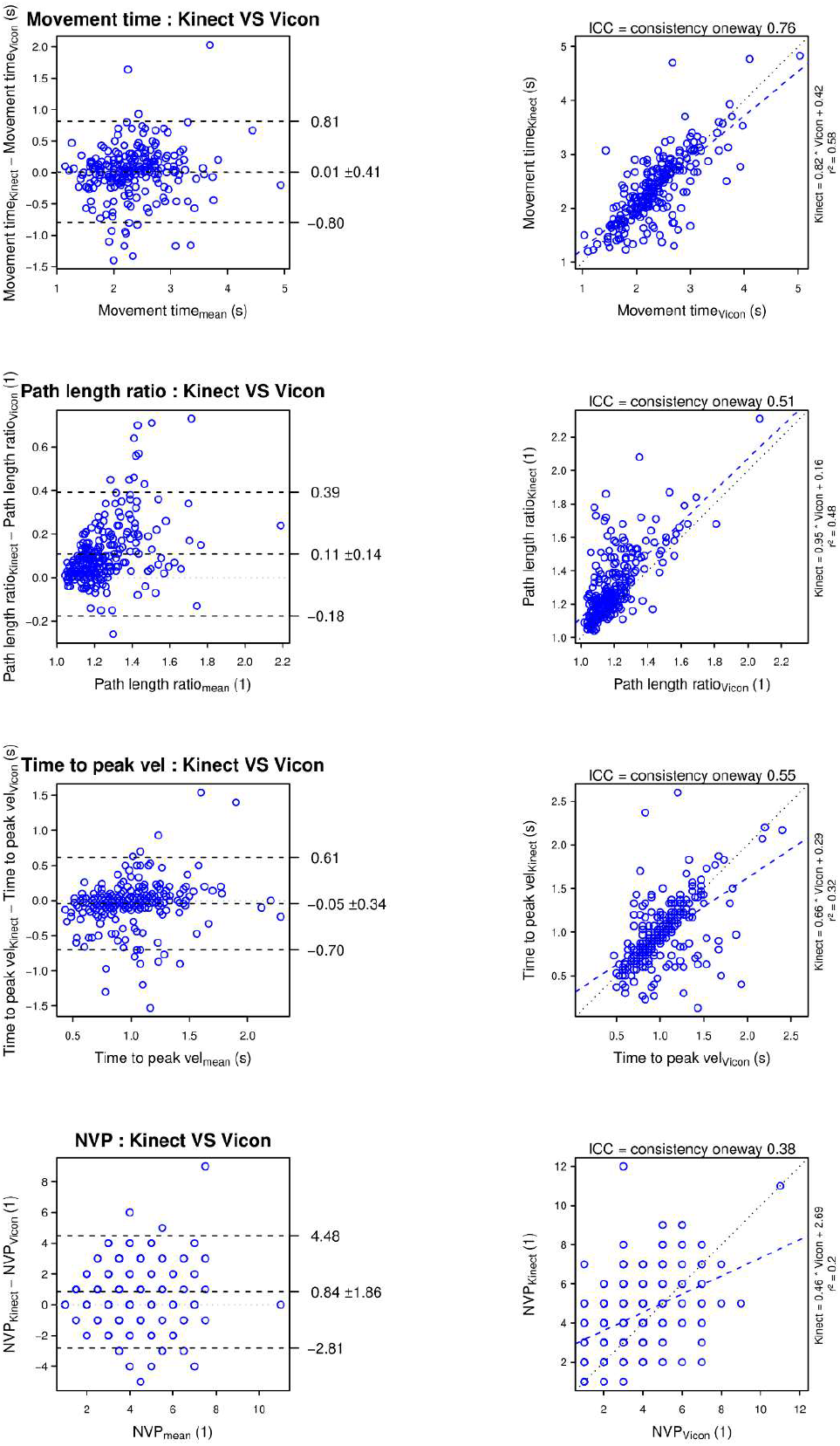
Comparison of hand efficiency, planning and smoothness assessed with the Kinect and the Vicon systems. Panels in the first row illustrate movement time (Movement time). Panels in the second row illustrate hand path length ratio (Path length ratio). Panels in the third row illustrate time to peak hand velocity (Time to peak vel). Panels in the fourth row illustrate number of hand velocity peaks (NVP). For each row, the left panel represents the Bland and Altman plot and the right panel represents the linear regression plot. When assessed with the Kinect, movement time is goodly reliable and have quasi no systematic error in average, path length ratio is moderately reliable and very slightly overestimated in average, time to peak velocity is moderately reliable and very slightly underestimated, number of velocity peaks is poorly reliable and slightly overestimated.

### 3.7. Speed

**Figure 5.**
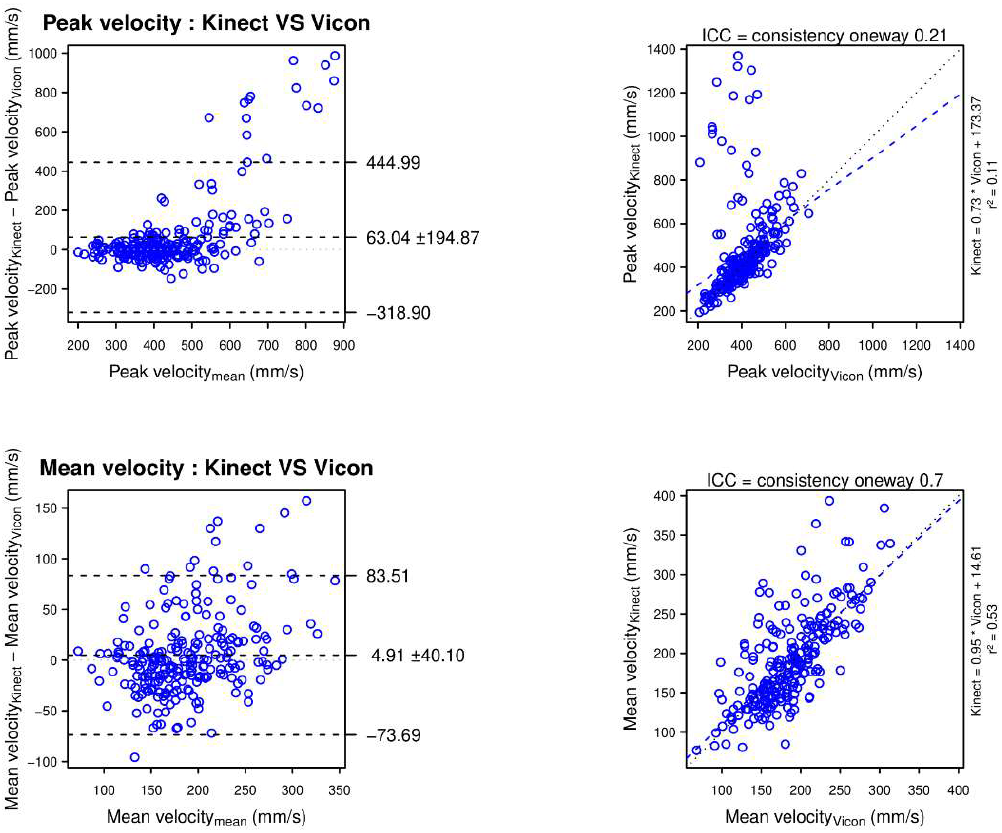
Comparison of the hand speed assessed with the Kinect and the Vicon systems. Panels in the first row illustrate peak hand velocity (Peak velocity). Panels in the second row illustrate mean hand velocity (Mean velocity). For each row, the left panel represents the Bland and Altman plot and the right panel represents the linear regression plot. When assessed with the Kinect, peak velocity is poorly reliable and slightly overestimated, mean velocity is moderately reliable and very slightly overestimated.

### 3.8. Displacements

The figure of displacements is in the supplementary data.

## 4) Discussion

Our study shows that in a horizontal reaching task, the Kinect measures trunk forward compensations with a good to excellent reliability and validity, but is unsensitive to low amplitude trunk rotation. The Kinect also measures hand and trunk range of motion as well as movement time and mean hand velocity with a moderate to good reliability, and with a good to excellent validity.

In contrast, the Kinect assesses variables involving elbow extension, shoulder flexion and shoulder abduction with a poor to moderate reliability and overall overestimates the variables. Finally, instantaneous Cartesian and angular measures with the Kinect are not precise enough which artificially creates jerky movements and overestimates NVP, Path Length Ratio, and Peak Velocity. Time to Peak Velocity is also affected resulting in a moderate reliability. Main results are summarised in Figure 6.

**Figure 6.**
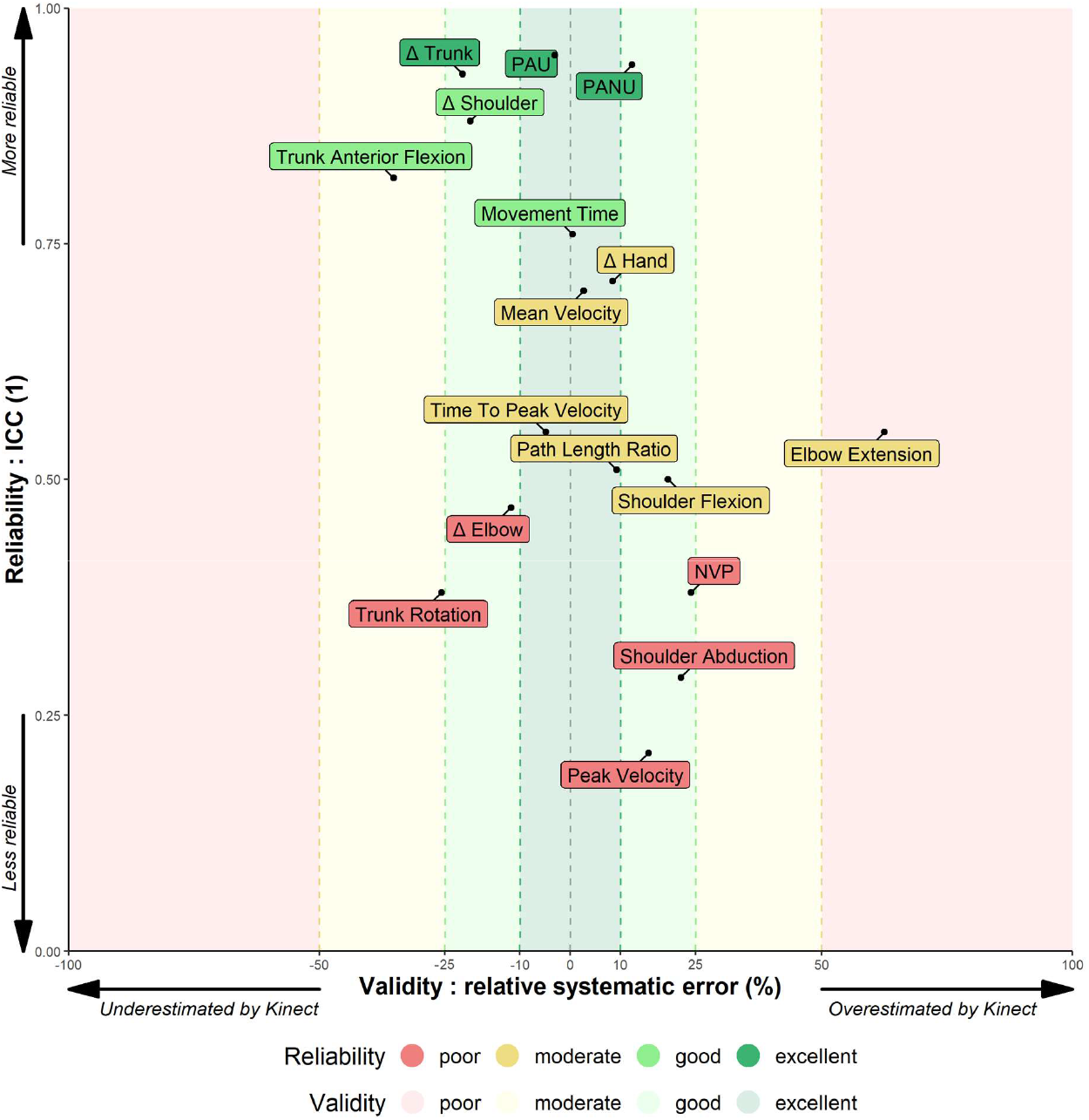
Summary of validity and reliability of 17 kinematic variables assessed by the Kinect. The X axis represents the validity through the relative error. The Y axis represents the reliability through the one-way ICC. The closer the variable is to the centre on the X axis, the more valid it is. The higher the variable is on the Y axis, the more reliable it is. A perfect match between the Kinect and the Vicon values would place the variable at the centre on the X axis (error = 0) and at the top on the Y axis (ICC = 1). The figure shows that averaged postural and angular measurements are much more reliable than instantaneous Cartesian measures.

These data indicate that, although the Kinect is not reliable enough to analyse fine kinematics over time, the Kinect does allow for global motion analysis (such as range of motion, movement time and mean velocity) with sufficient reliability in a clinical context.

### 4.1. Validation of the kinematic assessment obtained by Kinect

The aim of the study was to assess the reliability and validity of the Kinect main kinematic variables used in the analysis of reaching in stroke [38].

#### Trunk motion

The Kinect has excellent reliability on trunk displacement (ICC = .93) and good reliability on shoulder displacement (ICC = .88) and anterior flexion of the trunk (ICC =. .82). Because of a low mean trunk flexion (3.43°), the difference between means of −1.22 ± 3.05° overrepresent the error_relative_ (−35%) in the figure 6.

In contrast, trunk rotation was poorly assessed with the Kinect (ICC = .38), in particular with an underestimation of up to 20° of low trunk rotation (Fig. 3, top panel, left column) and other authors found the same result [18]. Note that the Vicon might also overestimate the low values of trunk rotation due to soft tissue artefact [39]. Indeed, trunk rotation being assessed mainly with the displacement of the shoulder marker on a transversal plane, a slight forward shift of the shoulder marker due to a shoulder flexion might artificially count for a trunk rotation. In any case, we propose that trunk rotation assessed with the Kinect should not be interpreted unless it exceeds 20°.

#### Hand motion

At the beginning of the reach, the Kinect sometimes confuses the forearm with the armrest. The Kinect correctly relocates the hand during the movement, creating a jerky correction. To a lesser extent, this temporo-spatial uncertainty occurs several times during the reach which artificially creates a jerkier movement, and overestimates the NVP and the path length ratio (Fig. 4, 2^nd^ and 4^th^ panel). By suddenly modifying the hand position, the Kinect creates a local velocity peak, possibly resulting in an overestimation of the peak hand velocity (Fig. 5, top panel). Due to its relation to peak velocity, the time to peak velocity is also affected in some cases (Fig. 4, 3^rd^ panel). By averaging the velocity over the entire movement, the mean velocity resists these local uncertainties (Fig. 5, bottom panel).

Because the reliability of Δ hand and Δ trunk assessment are good to excellent, variables derived from the Δ hand and Δ trunk measures such as PAU and PANU also have an excellent reliability, which confirms our previous findings [24].

#### The problem of elbow in a seated reaching task

First, the elbow is often confused with the back part of the armrest, resulting in a backward shift of the actual elbow position at the beginning of the reach. Second, due to the position of the hand located between the Kinect and the elbow, elbow occlusion can occur leading to an error in the elbow position. Third, elbow is poorly located on a fully extended arm (such as at the end of the reach) because of the alignment forearm - rupper arm. For these reasons, Δ elbow and side variables such as elbow extension, shoulder flexion and shoulder abduction are only poorly to moderately reliable and should be interpreted with caution.

To reduce elbow occlusion, some authors suggest installing the Kinect higher [18], in front of the participant [32, 33] and to perform functional movements in sitting [18, 40], but the present study shows that this is not enough to achieve sufficient precision for clinical interpretation of some variables. Other authors suggest installing the Kinect on the ipsilateral side of the movement with an angle of 30 to 45° [41], and moving the Kinect when assessing the other side, but this make the experiment more complicated and therefore reduces the fast and ease of use characteristic of the Kinect.

### 4.2. Smoothing out Kinect errors

The present study shows that the occlusion of the elbow and the confusion between the forearm and the armrest produce large errors resulting in a jerky movement. A first solution might be to use a chair with small armrests to decrease confusion errors. Furthermore, a correct filtering should be applied to the raw data. In fact, the analysis of raw data showed that the frequency band 0-2.5 Hz contains 95% of the spectral density. Applying a 2^nd^ order Butterworth type filtering with a cutoff frequency of 2.5 Hz greatly improved cartesian kinematics (table 2), while it had no effect on other variables (ICC changes ≤ .03). The ideal cut-off frequency for individuals with stroke might be higher than 2.5 Hz since people with stroke have more segmented movements than healthy individuals [42, 43]. Thus, a too low cut-off frequency might remove important information about the movement. However, our study shows that range of motion and averaged variables do not suffer from a lack of filtering, and can still be interpreted without filtering.

**Table 2.** Reliability improvements through filtering. The filtering consisted of applying a 2^nd^ order Butterworth filter with a cut-off frequency of 2.5 Hz on raw time series data. All variables shown in the table improved from filtering (ICC changes ≥ .12). Variables not shown in this table did not benefit from filtering (ICC changes ≤ .03).

|  | Movement time | Path length ratio | Time to peak velocity | NVP | Peak velocity |
| --- | --- | --- | --- | --- | --- |
| ICC before filtering | .64 | -.12 | .19 | -.30 | -.05 |
| ICC after filtering | .76 | .51 | .55 | .38 | .21 |

Other authors explored solutions to reduce Kinect errors. A deep learning algorithm applied to time series data reduced Kinect errors on shoulder and elbow range of motion by 88.8 ± 12.4 % [44]. Another approach is to fusion the data of several Kinects to minimise occlusions and optimise limb tracking [45–47]. Ryselis reported an overall increase in accuracy of 15.7%. A combination of a Kinect and several Inertial Measurement Units (IMUs) could also be used to reduce the upper limb position error up to 20% according to Jatesiktat *et al*. [48–50]. Finally, a device-independent approach is to incorporate body constraints (such as human skeleton length conservation and temporal constraints) to enhance the continuity of the estimated skeleton [51]. These solutions have the potential to consequently improve the accuracy of the Kinect while remaining affordable, even if the solutions require a high technical level and might lengthens the duration of patient preparation.

### 4.3. Kinematic assessment of the upper limb in clinical routine for personalised rehabilitation post-stroke

#### Markerless motion capture advantages for kinematic assessment

Due to its ease of use and markerless characteristic, the Kinect allows therapists to perform a simple kinematic assessment of the upper limb in about 5 minutes [3]. The software development kit (SDK) of the Kinect being openly available, the development of a software that automatically clean the data and compute the valuable kinematic variables is also possible. In addition, if the rehabilitation department does not already use markerless motion capture for virtual reality rehabilitation, the low cost of such devices ensures their accessibility.

#### Kinematic assessment to better understand the level of motor recovery of the patient

A seated reaching task with and without trunk restraint gives the therapist valuable information to better understand arm-trunk use [3], but the very same task, when recorded with the Kinect, opens the door to a more comprehensive kinematic assessment.

The movement time, the mean hand velocity, the path length ratio and the time to peak velocity reflect the spatial and temporal efficiency of the movement. A low mean hand velocity or a low time to peak velocity when combined to a high path length ratio or a high NVP reflect the increased importance of feedback and corrections during the reach and therefore signal a decreased efficiency of open-loop control [52, 53]. Except for the NVP, the Kinect measures these kinematic outcomes with acceptable accuracy-reliability (table 1), making the Kinect a valuable tool for monitoring changes in upper extremity kinematics over rehabilitation.

The elbow, shoulder and trunk range of motion describe the motor strategy and quantify the level of compensation used by the patient. Reduced elbow extension and shoulder flexion signal a deficit in upper limb movement that is often compensated by an increase of trunk flexion and trunk rotation, and a freeze of shoulder adduction [10, 54]. The Kinect measures these kinematic outcomes with varied accuracy-reliability making the Kinect a valuable tool for monitoring trunk flexion, but not for monitoring shoulder and elbow movements.

Quantifying the non-use of the shoulder-elbow joints with the PANU score [3] informs about what the patient can do, but does not do spontaneously. If the patient compensates with the trunk but can also perform the movement without using the trunk (PANU score > 6.5 %), then at least some of the compensation is not mandatory to succeed in the task. In contrast, if the patient has compensation and is unable to perform the movement without this compensation (PANU score < 6.5 %), then compensation is mandatory to succeed in the task [3]. This distinction is important because a compensation that improves reach efficiency should not be considered the same as a compensation which does not improve reach efficiency. The Kinect measures the PANU score with excellent accuracy-reliability (table 1), which confirms previous results [24].

#### Early and regular kinematic assessment to individualise rehabilitation, enhance motivation and improve recovery

Because kinematic monitoring is more sensitive to changes over the course of rehabilitation than clinical scales [6, 7], this could allow specific therapy to be planned based on specific changes in kinematics. For example, therapy focused on arm use makes sense for patients with high PANU scores but is certainly less suitable for patients with mandatory compensations.

Therapists can also provide kinematic feedback to patients and set goals to involve them in rehabilitation. Fine-tuned feedback enhances motivation and increases the level of acceptance of treatment by patients [55]. Thus, providing kinematic feedback to patients leads to better recovery [56, 57].

### 4.4. Limitations of the study

This study faces several limitations. The experiment was conducted on healthy volunteers whose movement characteristics were experimentally manipulated to approximate those of people who have suffered a stroke. Though it is reassuring that we here replicate the main results obtained with a Kinect and with patients [24], the Kinect-Vicon comparison with patients with stroke is a necessary additional step to go beyond the basic results presented here. Moreover, to induce a stroke-like movement, we asked healthy participants to hold a dumbbell during the reach. The presence of the dumbbell in the hand could have affected the ability of the Kinect to correctly locate the hand, even though this effect is likely small (i.e., the endpoint accuracy is similar for the loaded and unloaded hand and no significant difference was detected in the hand velocity root-mean-square error (RMSE_hand_) between conditions). The dumbbell could also have induced a greater occlusion of the elbow, which might have affected the performance of the elbow tracking, as evidenced by an increased error in the RMSE_elbow_ of 40 ± 72% between the unload and load conditions. However, this increased error in the elbow tracking is compensated by a wider dispersion in the load condition, and thus no improvement in the validity and reliability of the variables shown in this study was found in the unload condition. Based on the results presented here, it is effective to clinically assess upper limb kinematics after stroke with the Kinect v2, and this should be confirmed with a forthcoming clinical trial (https://clinicaltrials.gov/ct2/show/NCT04747587). Finally, the results presented here are only relevant for the same type of task, which is a unilateral horizontal seated reaching task. Results might differ in another type of task, such as non-horizontal reaching, or finger to noise test [58]. Indeed, due to the uncertainty of the depth measurement with the Kinect [15], movements in the frontal plane that are less dependent on depth changes might show a better accuracy than movements with variable depth when measured with the Kinect.

### 4.5. Future work

There is a need to replicate the validation in a population with movement disorders, whether for a Kinect or for any other markerless system with similar benefits. In addition, as Kinect-like systems are increasingly used in virtual reality rehabilitation to monitor the recovery process [13, 59], replication should focus on a wide variety of movements such as those used in virtual reality rehabilitation. Indeed, the present study assessed the validity of kinematic variables in a horizontal seated reaching task, but virtual reality protocols cover a much wider range of variables and tasks.

However, although Kinect is still widely used in virtual reality rehabilitation, Microsoft has stopped selling the Kinect since 2017. Instead, with the emergence of machine learning for image processing, many low-cost 2D joints tracking come to light [60], which led to 3D motion reconstruction modules based on multiple RGB cameras and joint triangulation as provided by OpenPose [61], VideoPose3D [62] or Learnable Triangulation [63]. Microsoft has also aligned itself with the release of the Kinect Azure in 2020, a new version of the Kinect that uses a deep neural network (DNN) method instead of the random forest method used by the Kinect v2 [64]. In order to anticipate their growing use, future work should assess the feasibility, validity and reliability of these systems in a clinical or home environment.

## 5) Conclusions

The aim of this study was to assess the validity and reliability of the Kinect v2 for quantifying upper body kinematics with application to monitoring upper limb function after stroke. Although therapists should be mindful of the Kinect limitations, particularly with respect to instantaneous kinematics, our results show that the Kinect quantifies reaching efficiency, compensation and nonuse with sufficient reliability to be used for rehabilitation purpose. The quantitative variables that are adequately monitored by the Kinect can effectively supplement the clinical scales used by therapists. In addition, a periodic assessment of the deficit can allow precise longitudinal follow-up of motor recovery, which could improve the evaluation of the rehabilitation modalities and help to optimise the therapeutic pathway of patients. A final advantage of using lightweight markerless motion capture devices in clinical routine is that an accumulation of kinematic data could allow ambitious retrospective studies to be carried out in the long run.

---

**<u>Supplementary figure 1 legend</u>**. *Comparison of nonuse and joint displacements assessed with the Kinect and the Vicon systems. Panels in the first row illustrate proximal arm nonuse (PANU). Panels in the second row illustrate proximal arm use (PAU). Panels in the third row illustrate trunk displacement (Δ Trunk). Panels in the fourth row illustrate shoulder displacement (Δ shoulder). Panels in the fifth row illustrate elbow displacement (Δ elbow). Panels in the sixth row illustrate hand displacement (Δ hand). For each row, the left panel represents the Bland and Altman plot and the right panel represents the linear regression plot. When assessed with the Kinect, PANU is excellently reliable but slightly overestimated, PAU is excellently reliable and very slightly underestimated, Δ trunk is excellently reliable and slightly underestimated, Δ shoulder is goodly reliable and slightly underestimated, Δ elbow is poorly reliable and slightly underestimated, Δ hand is moderately reliable and very slightly overestimated*.

## List of abbreviations

ICC: Intraclass Correlation Coefficient
NVP: Number of Velocity Peaks
PAU: Proximal Arm Use
PANU: Proximal Arm NonUse
RMSE: Root-Mean-Square Error

## Declarations

### Ethics approval and consent to participate

This study was performed in accordance with the 1964 Declaration of Helsinki. The Institutional Review Board of EuroMov at the university of Montpellier approved the study (IRB-EM 1901C).

### Consent for publication

Not applicable.

### Availability of data and materials

The code and the datasets generated and analysed during the current study are available in the Open Science Framework repository, https://osf.io/ckrdp/.

### Authors’ contributions

G.F. designed the protocol, recorded and analysed the data, wrote the first version of the manuscript. D.M. and J.F. assisted G.F. in designing the protocol, analysing the data and provided guidance for improving the manuscript. All authors accepted the latest version of the manuscript.

## Acknowledgements

We would like to thank Simon Pla for helping setting-up the experiment. We acknowledge Germain Faity for drawing the schema of the experimental conditions in Fig. 1.

