## Supplementary figures and images for "Validity and reliability of Kinect v2 for quantifying upper body kinematics during seated reaching"

### Supplementary figure 1

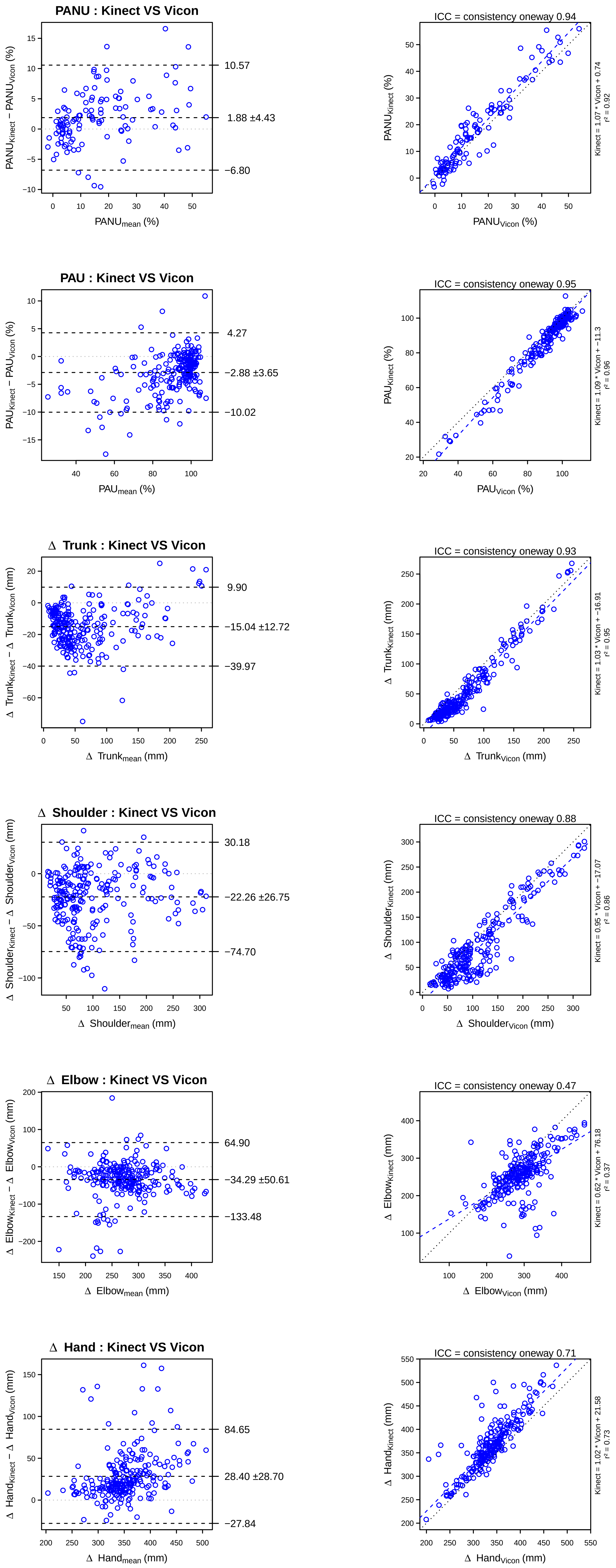
